# FGF12 Modulates Cardiomyocyte Hypertrophy by Inhibiting GATA4-ERK1/2 Phosphorylation Through Altered Intracellular Localization

**DOI:** 10.1101/2023.12.20.572709

**Authors:** Taojun Zhang, Kunlun Yin, Fangfang Yang, Tianjiao Li, Shuiyun Wang, Zhou Zhou

## Abstract

**Background:** Hypertrophic cardiomyopathy (HCM) is a heritable pathological condition resulting from mutations in sarcomere-related proteins, leading to severe structural abnormalities without effective treatment options. Although reduced *fibroblast growth factor 12*(*FGF12*) expression is observed in HCM patients, its functional role remains unclear.

**Methods:** Employing immunoprecipitation (IP)-mass spectrometry (MS) and CUT&Tag sequencing, we investigated FGF12-interacting proteins in myocardial samples from healthy volunteers and HCM patients. CRISPR-Cas9 was utilized to explore the function and interaction partners of FGF12 in cardiomyocytes induced from human pluripotent stem cells (hiPSCs-CMs), other cell lines, and mouse models (MYH7^R403Q^ and MYBPC3^c.790G>A^, transverse aortic constriction (TAC)). During hypertrophy, FGF12 localizes intranuclearly, prompting investigations into its binding to gene promoter regions through CUT&Tag sequencing and dual-luciferase experiments using myocardial tissues from patients. The beating frequency of hiPS-CMs was assessed using the CardioExcyte 96 real-time label-free cardiomyocyte functional analysis system.

**Results:** FGF12 was found to associate with proteins involved in energy metabolism, predominantly localizing to the perinuclear space under physiological conditions but shifting into the nucleus of hypertrophic cardiomyocytes. Co-IP-MS revealed significant interactions between FGF12 and metabolism-associated proteins, particularly GATA binding protein 4 (GATA4) and mitogen-activated protein kinase 1/3 (MAPK1/3) in the perinuclear space. In a hypertrophic state, FGF12 bound to the GATA4 promoter region, increasing its expression upon nuclear translocation. Both *in vitro* and *in vivo* models demonstrated that FGF12 interaction with GATA4 inhibited GATA4 and MAPK1/3 phosphorylation, inducing the expression of hypertrophy-associated genes. Overexpression of FGF12-NLS-del (nuclear localization signal deletion) resulted in decreased GATA4 phosphorylation, suggesting inhibition in the perinuclear region.

**Conclusions:** This study elucidates a pathological mechanism of HCM involving FGF12, where its nuclear localization enhances phosphorylation, GATA4 expression, and activation of the ERK1/2-pGATA4 pathway genes associated with hypertrophy. Beyond advancing our understanding of HCM, these findings propose FGF12 as a potential therapeutic target for HCM, warranting further exploration to potentially alleviate this condition affecting millions of individuals.

## Introduction

Hypertrophic cardiomyopathy (HCM) stands out as the most prevalent heritable heart disease, affecting approximately 1 in 200 individuals, with nearly 1500 reported mutations associated with the condition. The majority of these mutations manifest as structural abnormalities in sarcomere-associated proteins^1,2^, including thick myofilaments, sarcomere-binding proteins, troponin, and thin myofilaments, contributing to the formation of myocardial fibers^3^. Recent research has categorized hypertrophic cardiomyopathy models based on genotype(+/-) or phenotype(+/-), with G^-^/PH^+^ and G^+^/PH^+^ emerging as primary models^4^. Many reported mutations result in increased total energy production and ATP utilization, elevating myocardial energy requirements^5^. Consequently, inefficient energy utilization represents a prominent biophysical effect of myofilament mutations in HCM patients, accompanied by energy stress and adverse tissue remodeling^6^. To effectively treat HCM, a deeper understanding of methods to reduce myocardial energy levels is crucial.

*Fibroblast growth factor 12* (*FGF12*), a non-secreted protein belonging to the *fibroblast growth factor homologous factors (FHF)* family, is primarily expressed in human cardiac tissue^7^ and, in mice, in the brain^8^. Single-cell RNA-seq analysis has revealed a significant decrease in FGF12 expression in HCM patients compared to healthy individuals^9–11^. This reduction suggests a potential role for FGF12 in maintaining normal cardiomyocyte metabolism. Interestingly, FGF12 has been implicated in the regulation of cell cycle gene expression and the promotion of follicular granulosa cell proliferation through ERK phosphorylation in geese. However, the specific mechanism underlying these effects has remained elusive.

Two pivotal signaling pathways orchestrating cardiac hypertrophy are the NFAT (nuclear factor of activated T cells)/GATA4 pathway and the MAPK (mitogen-activated protein kinase)/ERK (extracellular signal-regulated kinase) pathway^12^. The ERK1/2 isoforms, collectively referred to as ERK, respond to stimulation by growth factors, displaying seemingly redundant functions. Upon activation, ERK translocates to the nucleus and phosphorylates multiple substrates^13^. Given the significance of cell growth and proliferation in heart development, the central role of ERK in cardiac physiology is anticipated^3^. Notably, ERK molecules have been implicated in various forms of cardiac hypertrophy and the progression to heart failure^14,15^. Despite the need for experimental characterization of the events leading to ERK activation, ERK or its MAPKs have been posited as a potential therapeutic target for treating cardiac hypertrophic diseases^16^.

The zinc finger DNA-binding transcription factor, GATA4 (GATA-binding protein 4), is well-established as an essential component for normal cardiac development and homeostasis in mice and humans. Mutations in GATA4 have been linked to heart defects in humans. ANP, BNP, cTnI, cTnT, and MYH7 are identified as downstream regulatory targets of GATA4. Among these, ANP and BNP serve as classical markers of hypertrophy in HCM^17^. Phosphorylation of the GATA4 residue S105 by ERK1/2 or MAPKs enhances its activity under cardiac stress. GATA4 activity is tightly associated with the downregulation of pro-fibrotic genes in the adaptive response to Ang-II. Additionally, the effects of GATA4 in cardiac fibroblasts hinge on S105 phosphorylation^18^. Despite this understanding, the role of p-GATA4 in the human myocardium is not fully elucidated, and it remains unclear whether and how it can be inhibited.

We generated three types of mouse models and MYH7 mutant induced pluripotent stem cardiomyocyte (iPS-CM) models of cardiac hypertrophy using CRISPR technology. Subsequently, we examined FGF12 at the sub-cellular level through staining human myocardial tissues and iPS-CMs. Our observations revealed its interaction with GATA4 and ERK1/2, as enriched by the pathway after coIP-MS in two normal controls and three HCM patients. To elucidate the role of FGF12 in different subcellar localizations, we assessed the changes in the aforementioned molecules following FGF12 full-length overexpression, FGF12-NLS-del (nuclear localization signal deletion) overexpression, and FGF12 knockdown. Given the nuclear localization characteristics of FGF12, we conducted CUT&Tag sequencing assays on myocardial tissues from patients with HCM. Notably, for the first time, we elucidated the biological role of FGF12 following its entry into the nucleus.

## Results

### FGF12 Expression is Decreased, and it re-localizes to the Nucleus in the Hypertrophic State

Prior research has indicated a substantial reduction in FGF12 expression in HCM patients with sarcomere mutations, along with altered subcellular localization during hypertrophic cellular conditions. In our previous histological analysis^7^, we observed a significant decline in FGF12 RNA expression in HCM patients carrying mutations in MYH7 or MYBPC3. To gain a more comprehensive understanding of FGF12 expression across various genetic backgrounds, we assessed RNA and protein levels of FGF12 in HCM patients with common mutations (G^+^/PH^+^), and those with nonspecific mutations (G^-^/PH^+^). While the G^-^/PH^+^ group exhibited consistently high expression, albeit not significantly different from healthy controls, the G^+^/PH^+^ HCM group showed a significant decrease in expression (Figure 1A, B, E, F). Additionally, FISH staining of heart tissues from both mutation groups revealed markedly lower FGF12 expression in G^+^/PH^+^ patients (Figure 1C, D). Notably, FGF12 quantification in mice demonstrated no statistical differences between the control group and the TAC, MYH7, and MYBPC3 groups (Supplemental 1B).

**Figure 1.**
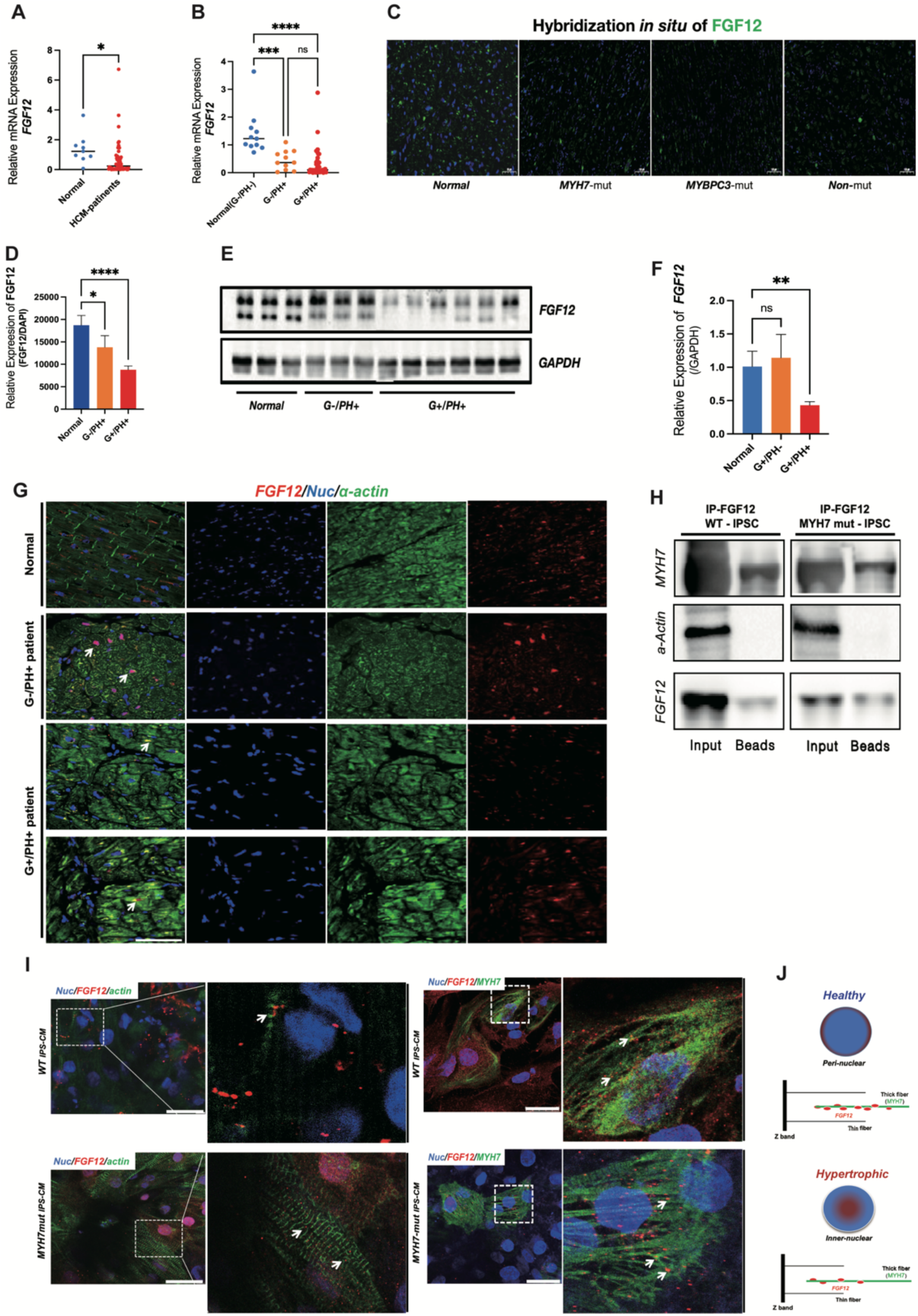
FGF12 co-localizes with thick myofilaments under normal physiological conditions and is re-localized to the nucleus under hypertrophic conditions. **A.** mRNA Expression of FGF12 in myocardial tissue comparing seven normal controls and 40 HCM patients, \**p* < 0.05. **B.** mRNA expression of FGF12 in myocardial tissues comparing 11 normal controls (genotype-/phenotype-), ten HCM patients (genotype-/phenotype+) versus 48 HCM patients (genotype+/phenotype+). \*\*\**p* < 0.005, \*\*\*\**p* < 0.001. **C, D**. FISH staining of myocardial tissue sections from normal control, MYH7-mut, MYBPC3-mut, and non-mut, where green represents FGF12 probe, blue represents DAPI. Fluorescence intensity counts the ratio of the average fluorescence intensity of ten 40× fields of view to the number of nuclei. *\*p <* 0.05, *\*\*\*\*p <* 0.001. **E, F.** Western Blot showing FGF12 expression in normal control, HCM patients (genotype-/phenotype+) vs. HCM patients (genotype+/phenotype+), where statistical values represent the grayscale logarithm of FGF12 bands vs. the endogenous gene GAPDH, *n. s* indicates no significance, *\*\*p <* 0.01. **G.** Immunofluorescence staining on paraffin sections of myocardium from normal control, HCM patients (genotype-/phenotype+) vs. HCM patients (genotype+/phenotype+); red represents FGF12, green represents α-actin, blue represents DAPI, and white arrowheads denote the co-localization of FGF12 and DAPI. **H.** Immunoprecipitation (IP) results of FGF12 in iPS-cardiomyocytes, FGF12 used as bait, IB antibody for MYH7 and α-actin; left panel for input group, right panel for the IP group. **I, J.** Immunofluorescence staining of FGF12 with α-actin and MYH7, respectively. FGF12 is shown in red in the upper panel, α-actin is shown in green in the lower panel, MYH7 is shown in green, and DAPI is shown in blue. The white arrow in the left panel indicates no co-localization between FGF12 and actin, while the white arrow in the right panel indicates co-localization between FGF12 and MYH7; J. is a schematic diagram of the localization of FGF12 with the nuclei, thick and thin myofilaments, with the upper panel depicting the localization of FGF12 in the normal control, and the lower panel illustrating it in the hypertrophic state.

Immunofluorescent staining of FGF12 protein in myocardial sections from patients (Figure 1G) and in cardiomyocytes differentiated from iPSCs (Figure 1I) validated the downregulation of FGF12, particularly in G^+^/PH^+^ cases. It also highlighted its predominant localization in the perinuclear region with co-localization with thick myofilaments in the physiological state. In contrast, in the hypertrophic state, FGF12 mainly localized in the nucleus with reduced co-localization with thick myofilaments (Figure 1J). Subsequent immunoprecipitation (IP) confirmed FGF12’s interaction with the thick myofilament marker, MYH7, but not the thin myofilament protein, α-actin. Moreover, interactions with MYH7 were diminished in the hypertrophied state (Figure 1H).

### FGF12 Overexpression Improves Myocardial Morphology, Reduces Heart rate, While Alleviating Cardiac Hypertrophy in Mice

In an effort to unravel the underlying mechanisms of FGF12-induced effects, we conducted IP coupled with qualitative mass spectrometry using samples from HCM patients with MYH7 point mutations and healthy volunteers. KEGG pathway enrichment analysis unveiled FGF12’s predominant involvement in energy metabolism-related pathways (Figure 2A). To delve deeper into FGF12’s role in energy metabolism, we assessed myocardial beating rhythm and amplitude in iPS-CMs using the CardioExcyte 96 real-time label-free cardiomyocyte functional analysis system. Notably, rhythm was lower in cardiomyocytes with FGF12 overexpression compared to negative controls (Figure 2B, C).

**Figure 2.**
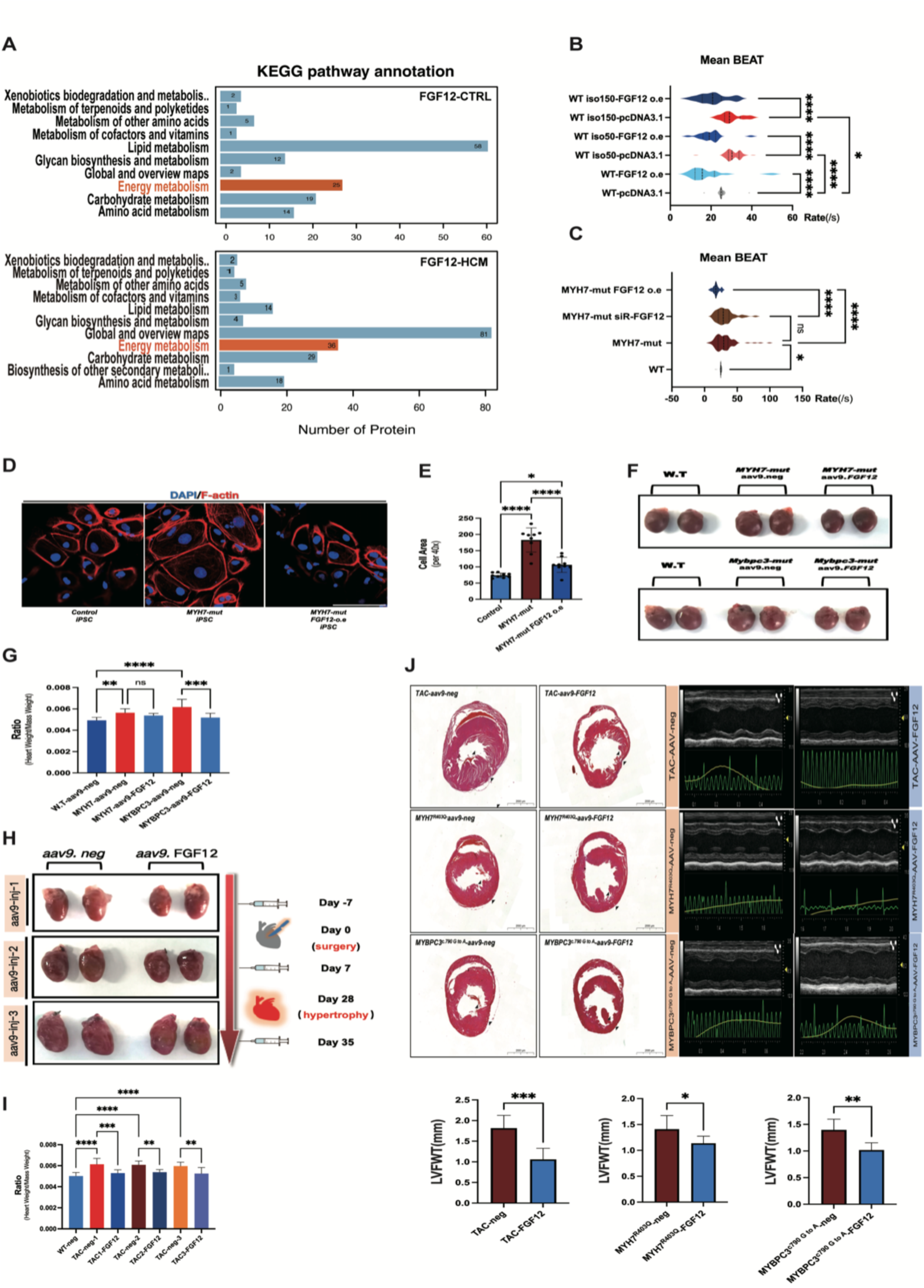
FGF12-mediated remodeling of myocardial morphology, energy metabolism, and alleviation of cardiac hypertrophy in various mice models. **A.** Differential expression plot of KEGG pathway enrichment by IP mass spectrometry in three MYH7-mut HCM patients and two normal controls. Red arrows indicate the concentration of FGF12-related proteins in the energy metabolism pathway, with the number of proteins shown on the horizontal axis. Energy metabolic pathways are labeled in red. **B, C.** CardioExcyte 96 real-time label-free cardiomyocyte functional analysis system illustrates the mean beats in different iPS-CMs. The horizontal axis represents the beating rates. Each group of cell samples ≧3, \**p* < 0.05, *\*\*\*\*p <* 0.001. **D, E.** Immunofluorescence staining shows cell morphology of normal, MYH7-mut, and MYH7-mut-FGF12 o.e., with F-actin in red and DAPI in blue. Cell area statistics represent the mean cell area in ten 40× fields of view, *\*p <* 0.05, *\*\*\*\*p <* 0.001. **F, G.** A schematic diagram of cardiac anatomy of *aav9-negative* and *aav9-FGF12* after injection in mice of different cardiac hypertrophy models (MYH7^R403Q^ and MYBPC3^c790^ ^G^ ^to^ ^A^). Plots depict the statistical results of heart-to-body weight ratio in different groups of mice, respectively, n=10, n.s. represents no significance, *\*\*p <* 0.01, *\*\*p <* 0.01, \*\*\**p* < 0.005, and *\*\*\*\*p <* 0.001. **H, I.** Schematic cardiac anatomy of aav9-negative and aav9-FGF12 injected TAC mice at different time points (1 week before surgery in the upper panel, 1 week after surgery in the middle panel, and 1 week after myocardial hypertrophy in the lower panel). Plots show the statistical results of the heart-to-body weight ratio of the different groups of mice subjected to TAC, respectively. n=10, *\*\*p <* 0.01, \*\*\**p* < 0.005, *\*\*\*\*p <* 0.001. **J.** Left panel: Cross-sectional H&E-stained images of different groups of mouse hearts, scale bar=2000 μm. Right panel: M-ultrasound echocardiography of the heart between aav-negative and aav-FGF12 group, histogram showing the difference in left ventricular free-wall thickness (LVFWT) between aav-negative and aav-FGF12 group. \*\**p* < 0.01, \*\**p* < 0.01, \*\*\**p* < 0.005.

Expanding on these findings, we overexpressed FGF12 in G^+^/PH^+^ iPS-derived cardiomyocytes and conducted immunofluorescence staining for F-actin to explore potential effects on size or morphology in hypertrophic cardiomyocytes. This analysis revealed that FGF12 overexpression led to significantly smaller G^+^/PH^+^ cells compared to those without overexpression (Figure 2D, E). Taken together, these results suggest that FGF12 overexpression can mitigate hypertrophic myocardial morphology, alleviate the heart rate, and reduce mitochondrial calcium concentrations.

Moving to *in vivo* experiments, we measured heart size and weight in transverse aortic constriction (TAC) model mice with or without heart-specific AAV9-FGF12 overexpression. AAV9-FGF12 or AAV9-negative-vector was injected at three different time points before and after TAC surgery. The results demonstrated variations in the heart/mass weight ratio among the three time points for each group, indicating that FHF12 may alleviate pressure-loading myocardial hypertrophy despite the absence of a statistical difference in weight (Figure 2H, I).

Building on these observations, we investigated whether FGF12 overexpression influenced heart size or weight in mice carrying the *MYH7^R^*^403^*^Q^* or *MYBPC3^c.^*^790^*^G>A^* mutations. In line with our previous findings, the cardiomyogenic ratio was lower in mutant mice with FGF12 overexpression compared to AAV9-negative controls, with no significant differences between the mutant groups (Figure 2F, G). H&E staining of myocardium sections from TAC or mutant mice revealed more pronounced thinning of the left ventricular free junction under FGF12 overexpression compared to vector controls (Figure 2J). The relief of myocardium following FGF12 overexpression suggests its potential to ameliorate myocardial hypertrophy by inhibiting specific hypertrophic factors.

### FGF12 Inhibits the Expression of Downstream Hypertrophy-related Genes by Inhibiting Phosphorylation of GATA4 (S105)

Initially, we scrutinized GATA4 levels in HCM patients. Western blot analysis of GATA4 and p-GATA4 (S105) levels in myocardial tissues revealed elevated levels in both G^+^/PH^+^ and G^-^/PH^+^ HCM patients compared to healthy controls (Figure 3A). Given that both FGF12 and GATA4 may exhibit nuclear or perinuclear localization, Co-IP experiments and staining were conducted to observe potential co-localization (Figure 3B, C), suggesting an interaction and co-localization between FGF12 and GATA4. Subsequently, iPS-cardiomyocytes were treated in different groups (DMSO-neg-vector, Iso-neg-vector, Iso-Si-FGF12, and Iso-FGF12-CDS). GATA4, p-GATA4 (ser105), and p-GATA4/GATA4 ratios were upregulated in the Isoproterenol-treated and DMSO groups. Transfection of Isoproterenol-treated iPS-cardiomyocytes with si-FGF12 FGF12-CDS plasmids resulted in a significant decrease in GATA4 and p-GATA4/GATA4 ratio in FGF12-CDS-iPS-cardiomyocytes (Figure 3D).

**Figure 3.**
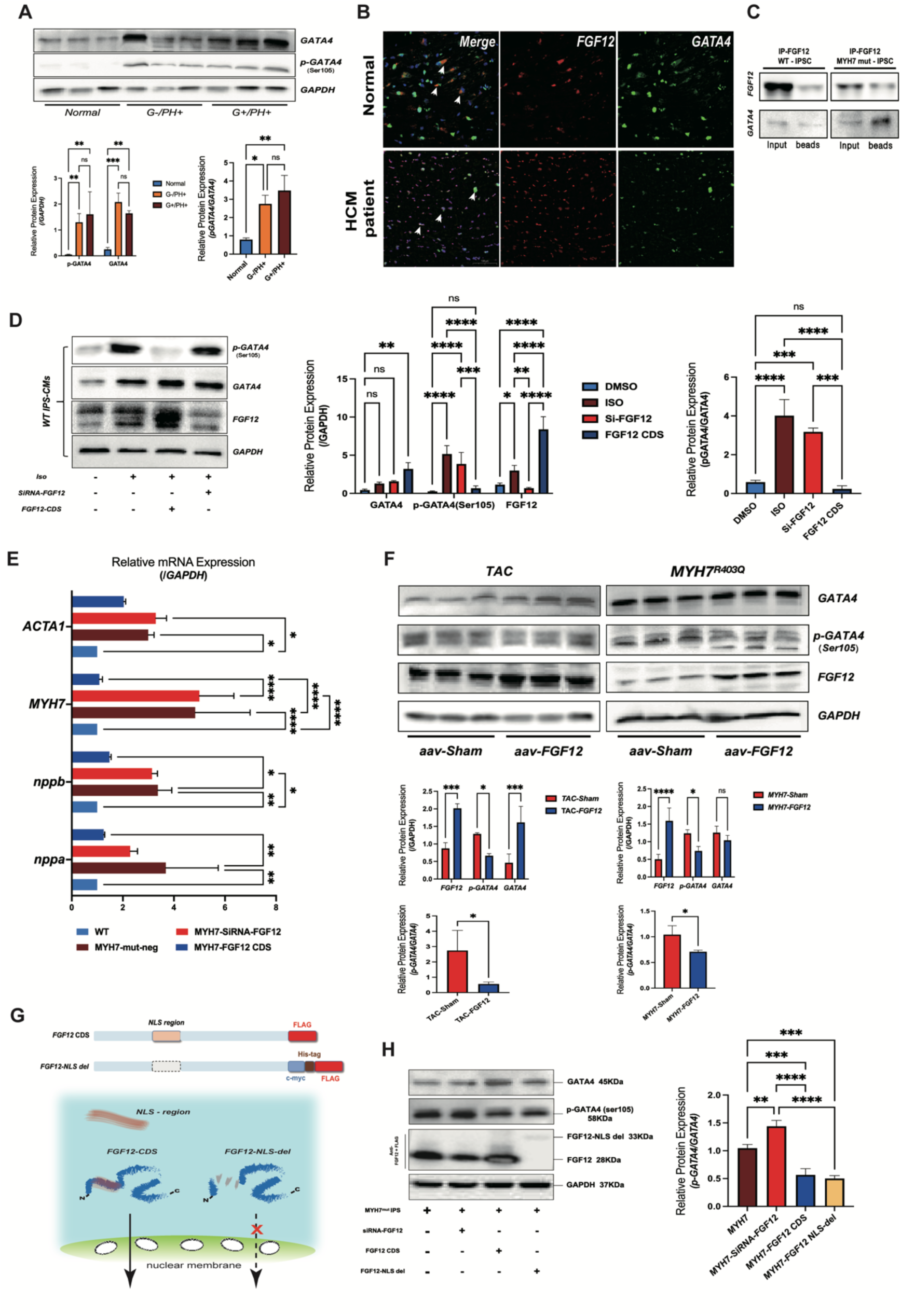
FGF12-mediated inhibition of GATA4 phosphorylation (S105) outside the nucleus. **A.** Protein expression results of p-GATA4(S105), GATA4, and GAPDH in three normal controls, three HCM patients (G-/PH+), and three HCM patients (G+/PH+) were statistically determined based on the relative expression of GATA4 and P-GATA4(S105) (normalized to GAPDH) and the ratio of P-GATA4(S105)/GATA4, \**p* < 0.05, \*\**p* < 0.01, *** *p* < 0.005. **B.** Immunofluorescence staining of normal control and HCM patient myocardium, with green for GATA4, red for FGF12, and blue for DAPI. White arrows indicate the co-localized region of FGF12 and GATA4, scale bar=100 μm. **C.** Results of immunoprecipitation experiments, IP antibody for FGF12, IB antibody for FGF12, and GATA4, respectively. Left panel is input, right panel is IP group. **D.** Western blot results of WT-iPS-CMs showing the expression results of p-GATA4(S105), GATA4, FGF12, and GAPDH in WT, isoprenaline, isoprenaline+siRNA-FGF12, and isoprenaline+FGF12-cds groups, respectively, and the statistics of the expression results of GATA4, P-GATA4(S105), and the FGF12 relative expression (normalized to GAPDH) and the ratio of P-GATA4/GATA4, *\*p <* 0.05, *\*\*p <* 0.01, *\*\*\*p <* 0.005, *\*\*\*\*p <* 0.001. **E.** q-PCR showing the statistical illustration of mRNA expression results of *nppa, nppb, acta1,* and *MYH7*, n≧3, \**p* < 0.05, *\*\*p <* 0.01, *\*\*\*p <* 0.005, *\*\*\*\*p <* 0.001. **F.** GATA4, P-GATA4, FGF12, and GAPDH expression in TAC mice and MYH7^R403Q^ mice. The statistics represent the relative expression of GATA4, P-GATA4 (S105), and FGF12 (normalized to GAPDH) and the ratio of P-GATA4/GATA4, *\*p <* 0.05, *\*\*\*p <* 0.005, *\*\*\*\*p <* 0.001. **G.** Schematic representation of the different plasmid constructs. The upper portion shows the linear structure of the plasmids, and the lower portion is a schematic representation of the localization of the various plasmids following entry into the cells. The FGF12-nls region is indicated in orange. **H.** The results of western blot after transfection of MYH7-mut iPS-CMs using different plasmids and siRNA-FGF12 with antibodies against FLAG, p-GATA4(S105), GATA4, FGF12, and GAPDH were statistically determined as the ratio of P-GATA4/GATA4, *\*\*p <* 0.01, *\*\*\*p <* 0.005, *\*\*\*\*p <* 0.001.

To further explore the effects of decreased p-GATA4 levels, qPCR analysis of p-GATA4-related downstream hypertrophy genes was performed, revealing suppressed expression in cells with high FGF12 expression (Figure 3E). Using TAC or MYH7^R403Q^ mice injected with AAV9-FGF12 or the vector control, the p-GATA4/GATA4 ratio was similarly reduced (Figure 3F).

To ascertain whether the effects of FGF12 were attributed to increased phosphorylation of GATA4 in the perinuclear region, we generated 3’FLAG-tagged FGF12 variants with or without the nuclear localization signal (FGF12-CDS-FLAG and FGF12-NLS-del-FLAG, respectively, Figure 3G). Plasmids with these constructs or siRNA-FGF12-GFP/siRNA-NC-GFP were transfected into MYH7 mut IPS-CMs. Western blot analysis indicated decreased p-GATA4 levels and p-GATA4/GATA4 ratio in cells expressing FGF12-CDS-FLAG or FGF12-NLS-del-FLAG relative to the vector control and FGF12 knockdown IPS-CMs (Figure 3H). These results suggest that FGF12 can inhibit the phosphorylation of GATA4 in the perinuclear area, thus suppressing the downstream expression of hypertrophic genes. Notably, we observed an overall increase in the expression level of GATA4 in cells and mice after full-length overexpression of FGF12, indicating a potential role in the nucleus.

### FGF12 Entry into The Nucleus Promotes the Upregulation of GATA4 Through Binding to its Promotor Region

CUT&Tag analysis demonstrated the enrichment of FGF12 in the GATA4 promoter region. A query of the UCSC database (University of California, Santa Cruz, https://genome.ucsc.edu) for potential recognition/binding motifs in the GATA4 promoter sequence unveiled that FGF12 could potentially bind to GATA motifs located at positions GATAAG and AGACAA in the GATA4 promoter (Figure 4A, B). To validate whether FGF12 could interact with the GATA4 promoter to upregulate its expression, luciferase assays were conducted in HEK-293A cells expressing FGF12 and FGF12-NLS-del with the GATA4 promoter as bait (Figure 4B, C) The results indicated that the GATA4 promoter was indeed activated in the presence of FGF12 but not in the absence of the FGF12-NLS, signifying that FGF12 nuclear localization was required for its interaction with the GATA4 promoter and the promotion of GATA4 expression in cardiomyocytes.

**Figure 4.**
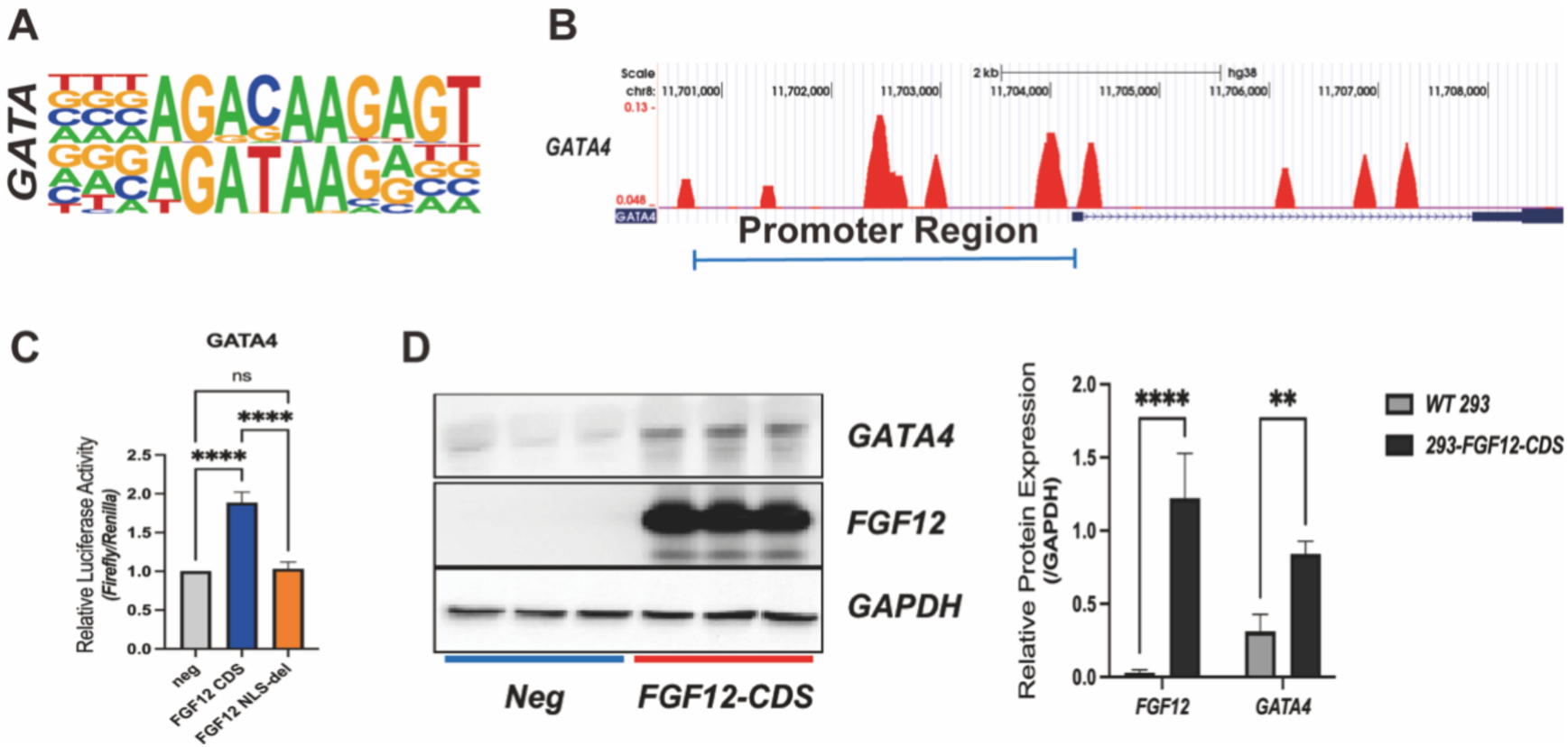
Intranuclear FGF12 interaction with the GATA4 promoter enhances GATA4 expression. **A.** CUT&Tag results indicating FGF12 enrichment in the motif plot of GATA4, represented as AGACAAGAGT. **B.** Schematic diagram illustrating FGF12 binding to the promoter region of GATA4 using UCSC. The red wave represents the number of reads, and the blue scale shows the promoter region of GATA. The blue bar indicates the length of GATA4’s promoter, scale bar=2kb. **C.** Dual-luciferase results displaying relative fluorescence intensity values (firefly/Renilla) in HEK293 cells transfected with FGF12, FGF12-nls-del, and an empty plasmid, respectively, *\*\*\*\*p <* 0.001. **D, E.** Quantification of GATA4 protein after transfection of HEK293 cells with negative and FGF12-cds plasmids. The histogram illustrates the grayscale statistics of the bands (normalized to GAPDH), *\*\*p <* 0.01, *\*\*\*\*p <* 0.001.

### FGF12 Inhibits the Phosphorylation of ERK1/2

GATA4 phosphorylation by ERK has been demonstrated to enhance GATA4-DNA binding and activate the hypertrophic gene program^19^. Given that FGF12 has previously been shown to inhibit the phosphorylation of ERK1/2 in follicular granulosa cells^20^, we sought to confirm the relevance of these findings in cardiomyocytes. Building upon our earlier data indicating a reduction in beating rate and amplitude of hypertrophic IPS-CMs under FGF12 overexpression, and mass spectrometry data revealing enrichment for the energy metabolism pathway (especially MAPK1/3, Supplemental 2F) among FGF12-binding proteins, we hypothesized that FGF12 could potentially reduce energy metabolism in hypertrophic cardiomyocytes by inhibiting the MAPK/ERK pathway. Western blot analysis of clinical samples showed that p-ERK1/2 levels were significantly higher in HCM patients than in healthy controls (Figure 5A, B). Similarly, the p-ERK1/2 levels and the p-ERK1/2/ERK1/2 ratio were significantly lower in TAC and MYH7^R403Q^ model mice overexpressing FGF12 compared to the vector control mice *in vivo* (Figure 5C, D). We also examined p-ERK1/2 and ERK1/2 levels under FGF12 silencing or overexpression in WT iPS-CMs treated with Iso or MYH7-mut iPS-CMs and found that p-ERK1/2 levels and the p-ERK1/2/ERK1/2 ratio were both reduced in cells overexpressing *FGF12* (Figure 5E, F). These consistent results among murine and human samples *in vitro* and *in vivo* support the likelihood that FGF12 could regulate the phosphorylation of GATA4 via modulation of p-ERK1/2 levels.

**Figure 5.**
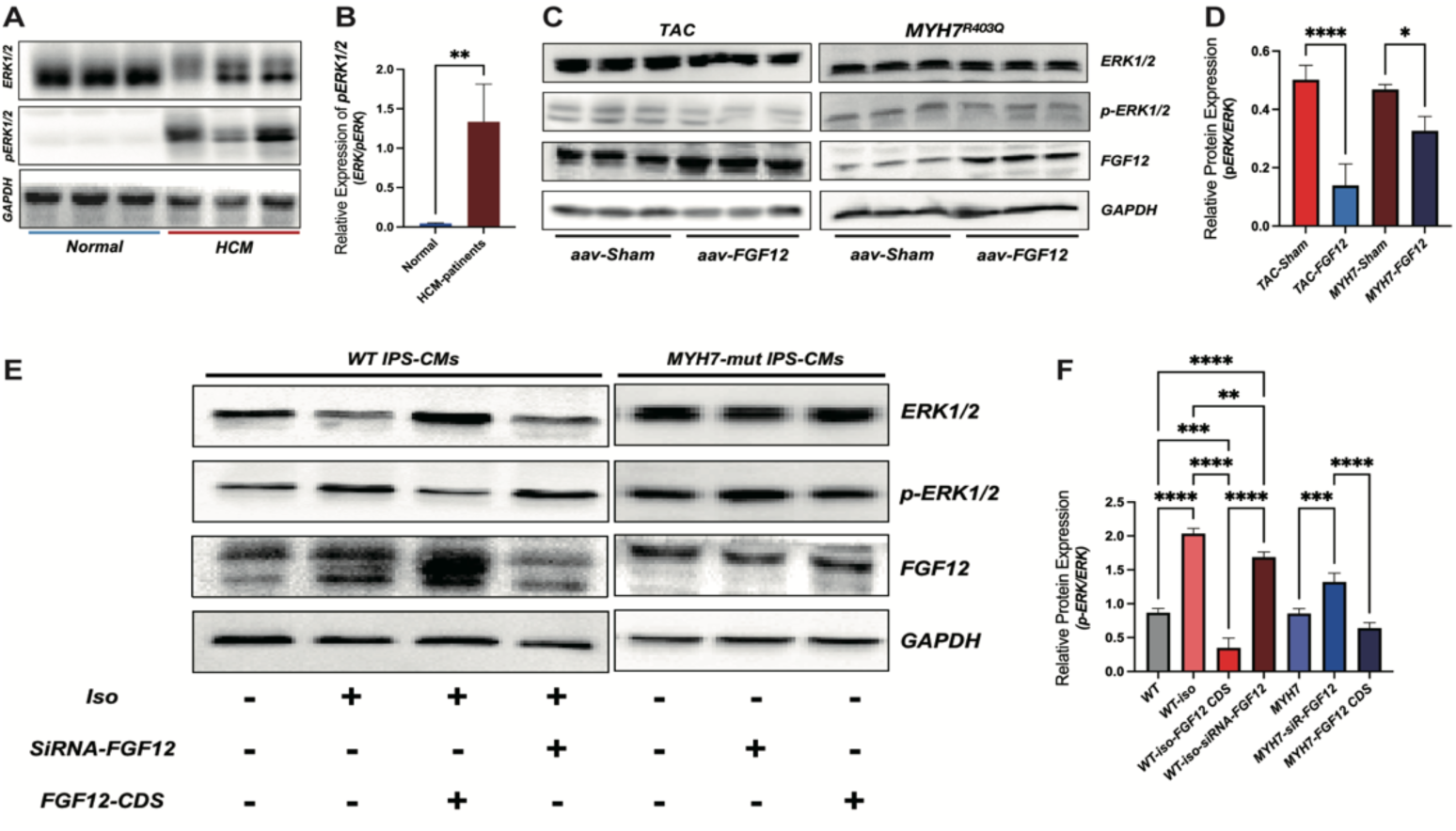
FGF12 suppresses phosphorylation of ERK1/2. **A, B.** Protein quantification results of three normal controls and three HCM patients using antibodies against ERK1/2, P-ERK1/2, and GAPDH. The statistics include the ratio of P-ERK1/2**/**ERK1/2, *\*\*p <* 0.01. **C, D.** Western blot results of TAC and MYH7^R403Q^ mice with antibodies against ERK1/2, P-ERK1/2, and GAPDH. The statistics represent the ratio of P-ERK1/2**/**ERK1/2 in various groups, *\*\*p <* 0.01, *\*\*\*\*p <* 0.001. **E, F.** Protein quantification results of three normal controls and three HCM patients with antibodies against ERK1/2, P-ERK1/2, and GAPDH. The statistics include the ratio of P-ERK1/2**/**ERK1/2 in various groups, \*\**p* < 0.01, *\*\*\*p <* 0.005, *\*\*\*\*p <* 0.001.

## Discussion

Building upon our staining results, which demonstrated that FGF12 localizes with the myocardial thick myofilament marker MYH7 and co-localizes with mitochondria within human myocardial tissue, we hypothesize that FGF12 might interact with MCU (Mitochondrial Calcium Uniporter) on mitochondria (Supplemental 1C). Subsequent co-IP-MS results further suggested that FGF12 could interact with proteins related to cardiomyocyte metabolism. As induced pluripotent stem cardiomyocytes (iPS-CMs) closely mimic human cardiomyocytes *in vitro*, we validated our staining results in iPS-CMs, supporting observable differences in localization and function of cardiomyocytes in distinct physiological and pathological states.

To recapitulate the pathogenic process of human hypertrophic cardiomyopathy *in vitro*, we utilized CRISPR-Cas9 to construct an MYH7 c.1988G>A: p.Arg633His thick myofilament mutant with pathogenic grade P, along with an isoproterenol-induced hiPS-CM cell line. Additionally, we generated another iPS-CM line harboring a thin myofilament mutation (*ACTC1* c.715G>C p.Glu239Gln) (strategy in Supplemental 1A). Given that FGF12 knockdown does not result in a clear phenotype *in vivo* due to its extremely low expression in murine heart tissues^21^, we simulated hypertrophic cardiomyopathy of different etiologies in mice, such as the MYH7-mut (R403Q) and MYBPC3-mut (c.790 G to A) lines, and a surgery-induced TAC model. In all three models, we observed a significant improvement in myocardial hypertrophy within one week after the AAV9-FGF12 injection, most notably in the left ventricular free wall. While these results suggest that FGF12 conferred some therapeutic effects in mice with myocardial hypertrophy, we also noted that fibrosis was exacerbated after myocardial hypertrophy in mice treated with FGF12 (Supplemental 2A-D), consistent with the findings of previous studies in liver fibrosis^21^.

Our findings highlight FGF12 as a highly conserved gene, as evidenced by our attempts to generate a homozygous FGF12^-/-^ IPS line through CRISPR-Cas9 in IPSCs prior to CM induction (Supplemental 1A). This unattainability persisted in HEK-293 cells and mesenchymal stem cells, underscoring the indispensable role of FGF12 in human cells. Quantitative assessment of FGF12 protein in human tissues unveiled the presence of two distinct FGF12 bands, even with different antibodies. We postulated that these bands might originate from different intranuclear and extranuclear FGF12 transcripts, aligning with our tissue staining results indicating intranuclear localization. However, nucleoplasm isolation with protein quantification in various iPS-CM lines revealed the presence of both protein bands and, consequently, both transcript isoforms inside and outside the nucleus. IP-MS results indicated that the subcellular localization proportion of FGF12-interacting proteins was generally consistent in the two groups, with slight variations from the staining results (Supplemental 1D).

It is noteworthy that, in mice, FGF12 expression is confined to brain tissue, with extremely low levels in cardiac tissue. This observation aligns with previous reports that found no discernible effect on cardiac phenotype in mice with systemic FGF12 knockout. Single-cell RNA-sequencing (ScRNA-seq) data from our HCM patients indicated the most significant decrease in FGF12 expression, temporally corresponding with the upregulation of *nppb*, a pivotal indicator of cardiac function^9^. Although FGF12 is a non-secretory protein undetectable in the peripheral blood, its expression generally correlates positively with cardiac function. A more pronounced decline in FGF12 expression is associated with a greater decline in cardiac function, prompting our exploration of whether exogenous FGF12 treatment could reverse hypertrophic cardiomyopathy in mice.

GATA4 is a well-established essential transcription factor in cardiac development, with its expression significantly increased during hypertrophic cardiomyopathy. Under normal physiological conditions, GATA4 is distributed in both perinuclear and intranuclear regions. However, during myocardial hypertrophy, GATA4 undergoes phosphorylation and strictly localizes in the nucleus^21–23^. Our results indicate that GATA4 co-localizes with FGF12 in perinuclear regions under physiological conditions and shifts to the nucleus in hypertrophic cardiomyocytes. It remains unclear whether the nuclear translocation of FGF12 coincides with GATA4 entering the nucleus. In cells overexpressing FGF12 or FGF12-NLS del, the levels of p-GATA4 (ser105) decrease, suggesting that the perinuclear localization of FGF12 may function to inhibit the levels of phosphorylated GATA4 in the nucleus, thereby limiting the expression of hypertrophy-related genes.

Conversely, in the disease state, FGF12 translocating into the nucleus results in decreased binding and increased phosphorylation of perinuclear GATA4. The subsequent entry of p-GATA4 into the nucleus then upregulates hypertrophy-related genes. This increased ratio of nuclear p-GATA4 and FGF12 to perinuclear GATA4 likely forms a pathogenic positive feedback loop regulating hypertrophy in cardiomyocytes. Additionally, extranuclear FGF12 inhibits ERK phosphorylation, while depleting perinuclear FGF12 by translocating to the nucleus promotes ERK1/2 phosphorylation, ultimately leading to myocardial hypertrophy (Figure 6).

**Figure 6.**
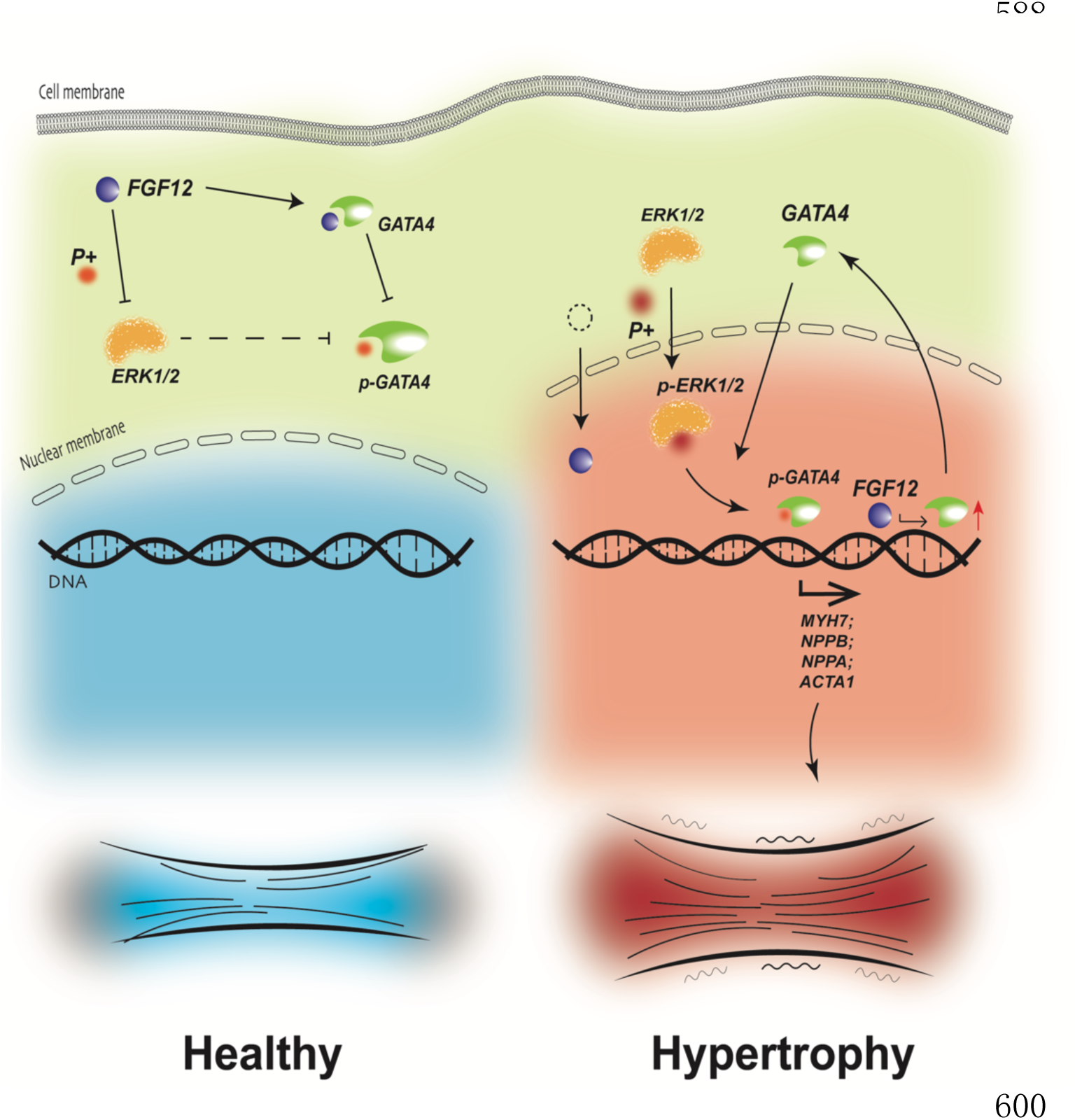
FGF12 inhibits cardiac hypertrophy when localized in the perinuclear area and promotes cardiac hypertrophy when entering the nucleus. In the normal physiological state, FGF12 is primarily located in the perinuclear region, where it inhibits the phosphorylation of ERK1/2, consequently suppressing downstream phosphorylation of GATA4. This inhibitory mechanism helps maintain normal cardiac function and morphology. In the hypertrophic state, the reduced presence of FGF12 in the perinuclear region leads to an increased level of phosphorylation of ERK1/2 in the perinuclear area. This enhanced phosphorylation of ERK1/2 induces the phosphorylation of GATA4, resulting in the elevated expression of hypertrophic genes (*nppa, nppb, MYH7,* and *ACTC1*). Simultaneously, the increased nuclear localization of FGF12 contributes to the heightened expression of GATA4, forming a feedback loop involving GATA4-ERK1/2-p-GATA4. This loop amplifies hypertrophy in cardiomyocytes, exacerbating the hypertrophic state.

In summary, the perinuclear localization of FGF12 can inhibit cardiac hypertrophy by inhibiting GATA4-ERK1/2 phosphorylation. Conversely, nuclear localization of FGF12 increases p-GATA4, amplifies GATA4 expression, activates the pathological expression of hypertrophy-associated genes, and consequently leads to cardiac hypertrophy.

---

**Table S1.**
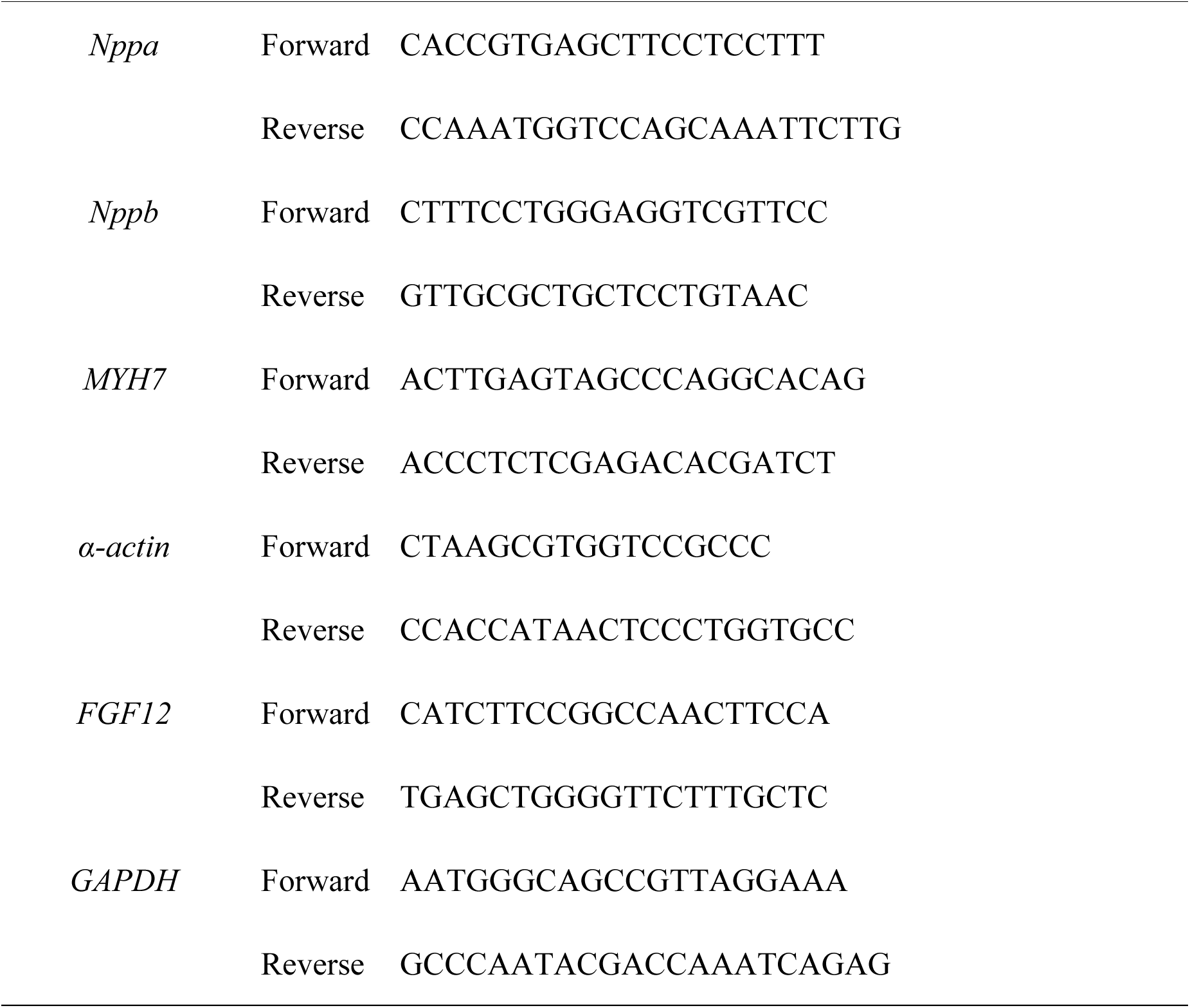
q-PCR primers.

**Table S2.**
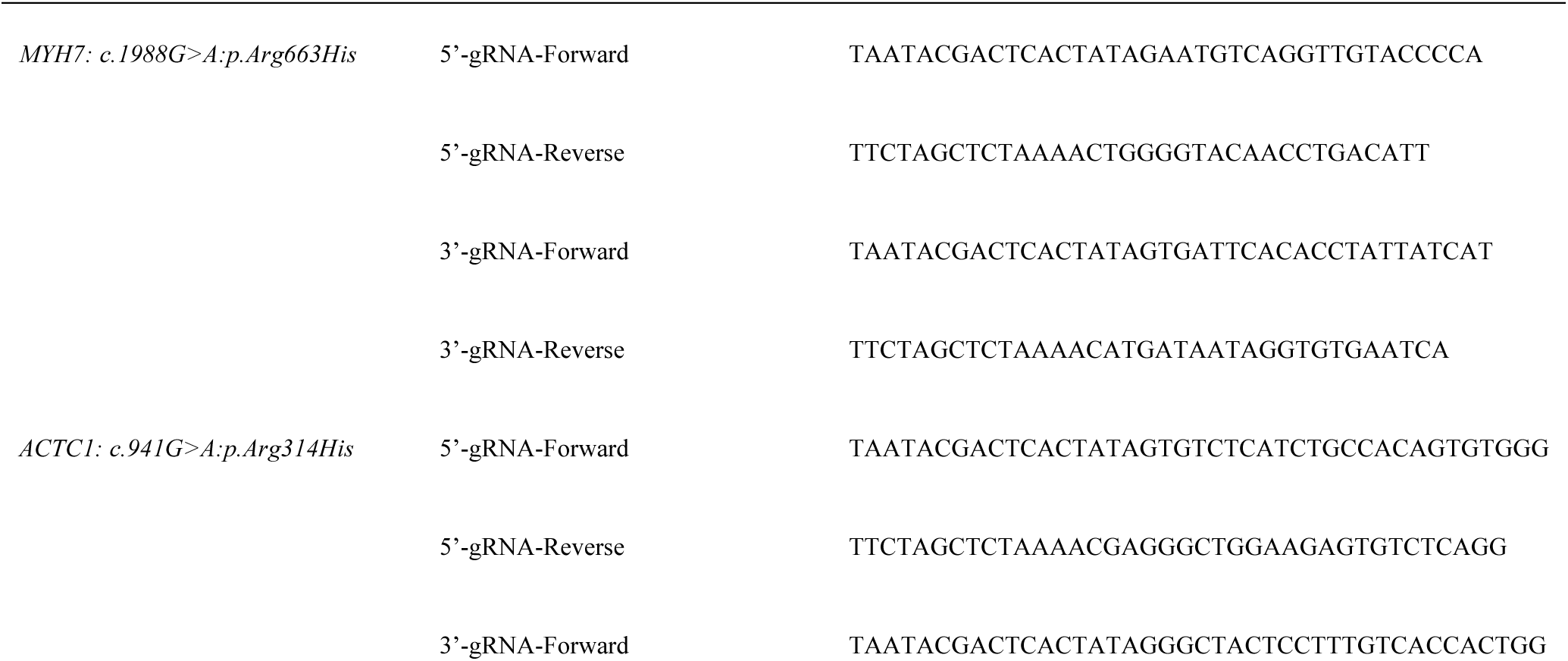
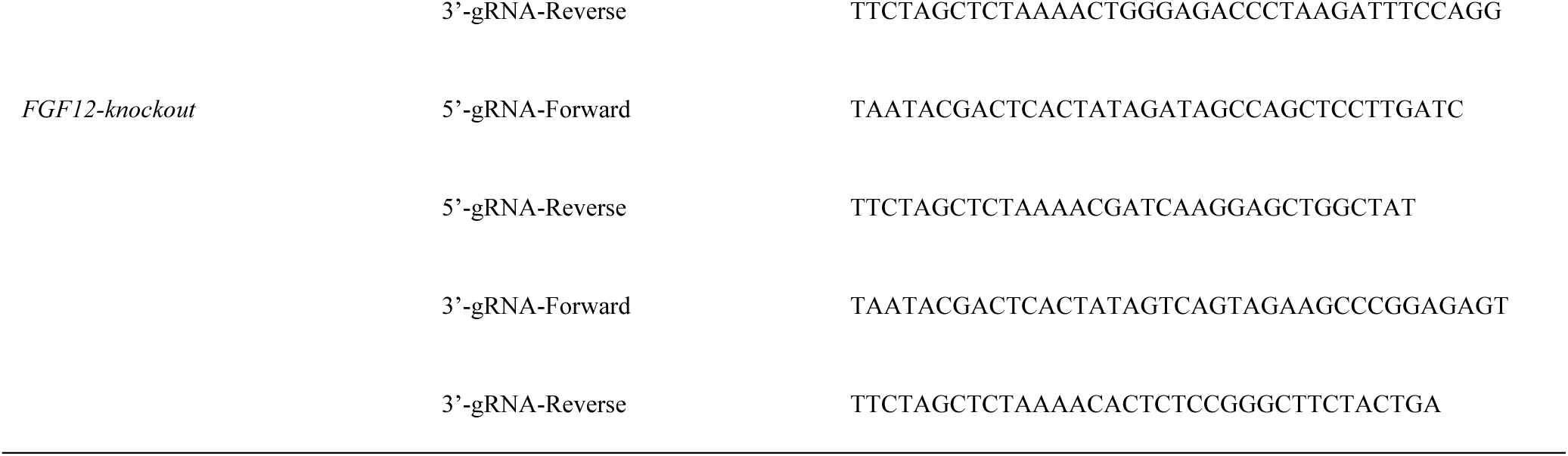
Template gRNA primers & identified.

**Table S3.**
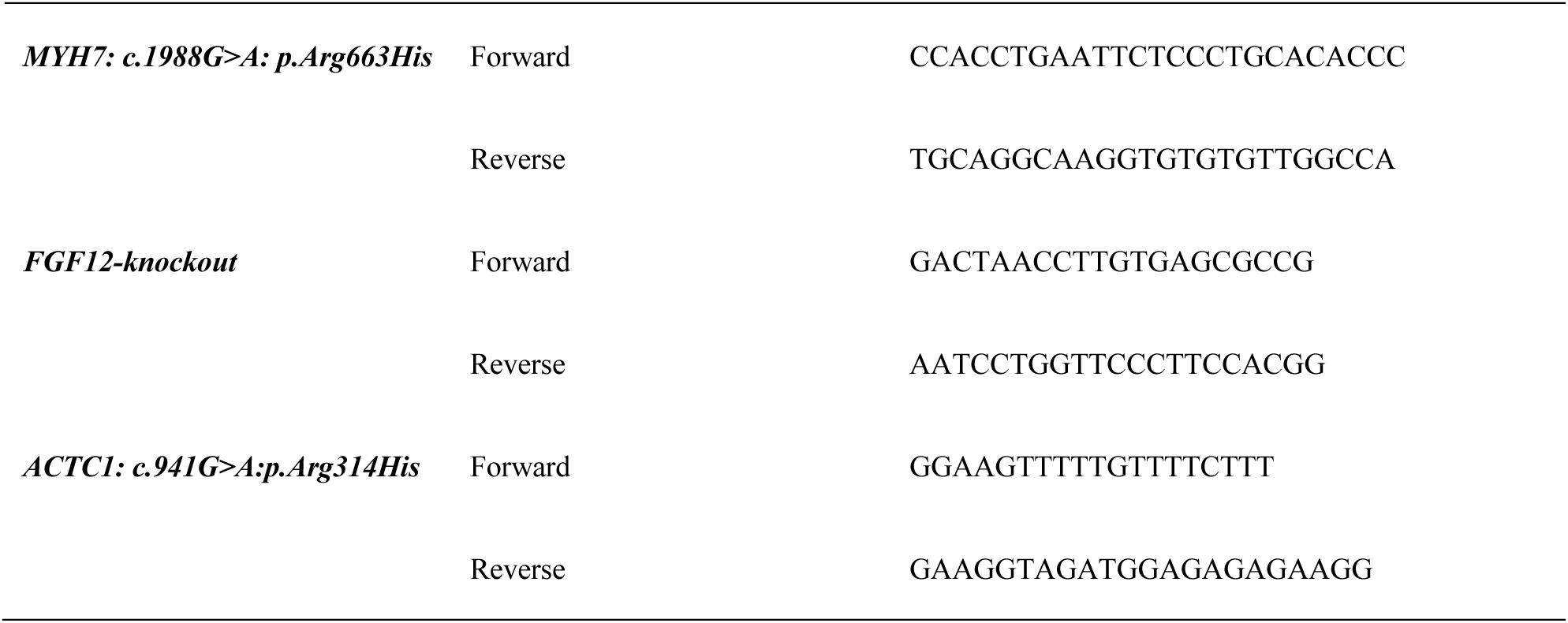
Primers for Identification.

**Supplemental Figure 1.**
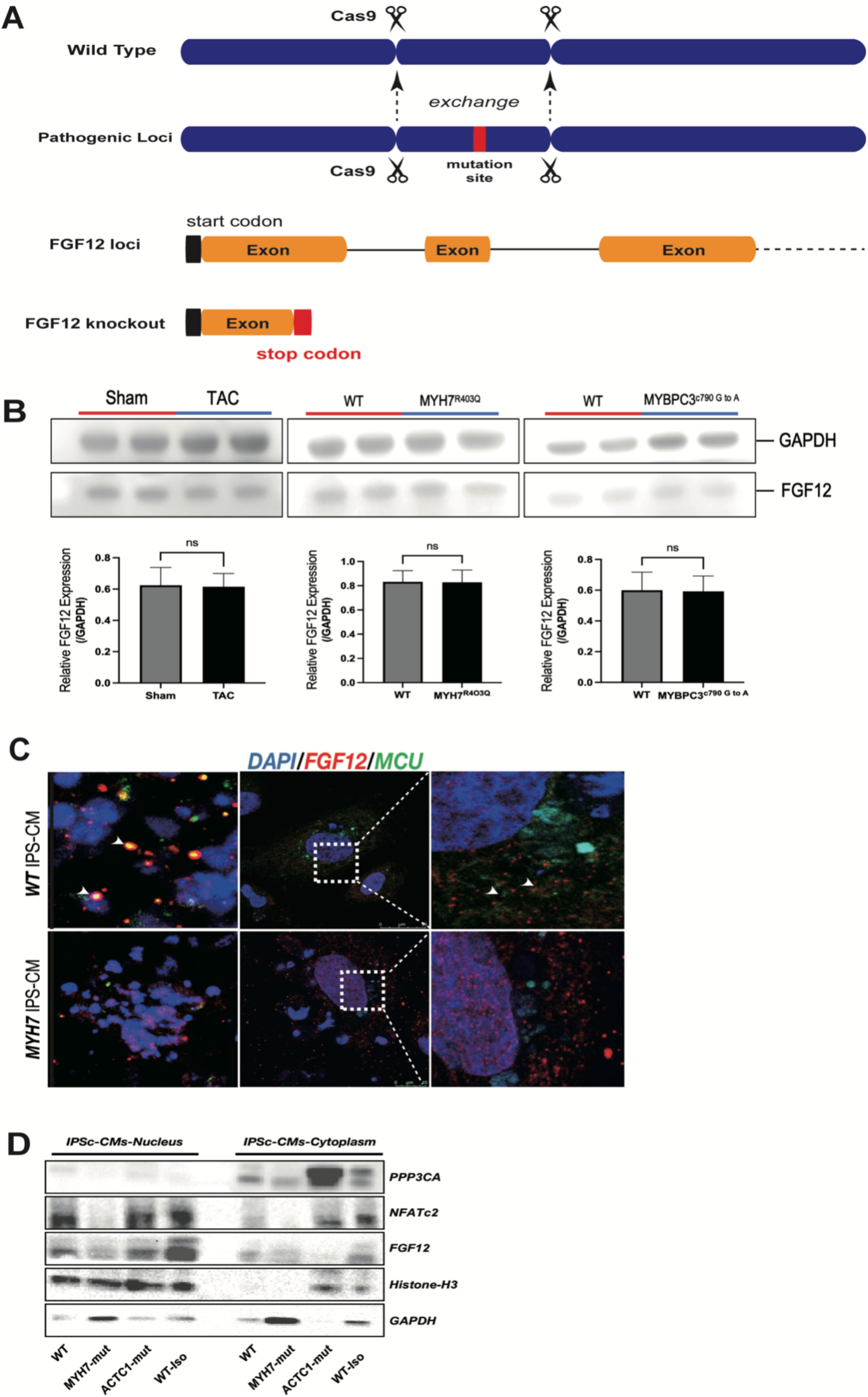
**A.** CRISPR-Cas9 strategy for pathogenic modeling and FGF12 knockout in iPSc lines. Upper panel: Cas9 protein cleaves homologous sequences in mutated and normal regions, resulting in the introduction of the disease mutation. Lower panel: Insertion of a stop codon TGA in the exon region of FGF12 leads to the knockout of FGF12 in iPSc lines. **B.** Western blot results of TAC MYH7^R403Q^ and MYBPC3^c790^ ^G^ ^to^ ^A^ mice using antibodies against FGF12 and GAPDH. The statistical evaluation of the relative expression of FGF12 across various groups, n.s., represents not statistically significant. **C.** Immunofluorescence co-staining results of FGF12 with MCU. Top panel: with WT-iPS-CMs. Bottom panel: with MYH7mut-iPS-CMs. Arrows indicate co-staining areas of FGF12 with MCU, with a scale bar of 25 μm. **D.** Nucleo-plasmic isolation assay of iPS-CMs. Left panel: separation of nuclear and plasma proteins, with antibodies against PPP3cA, NFATc2, FGF12, GAPDH, and histone-H3. Right panel: separation of cytoplasmic proteins.

**Supplemental Figure 2.**
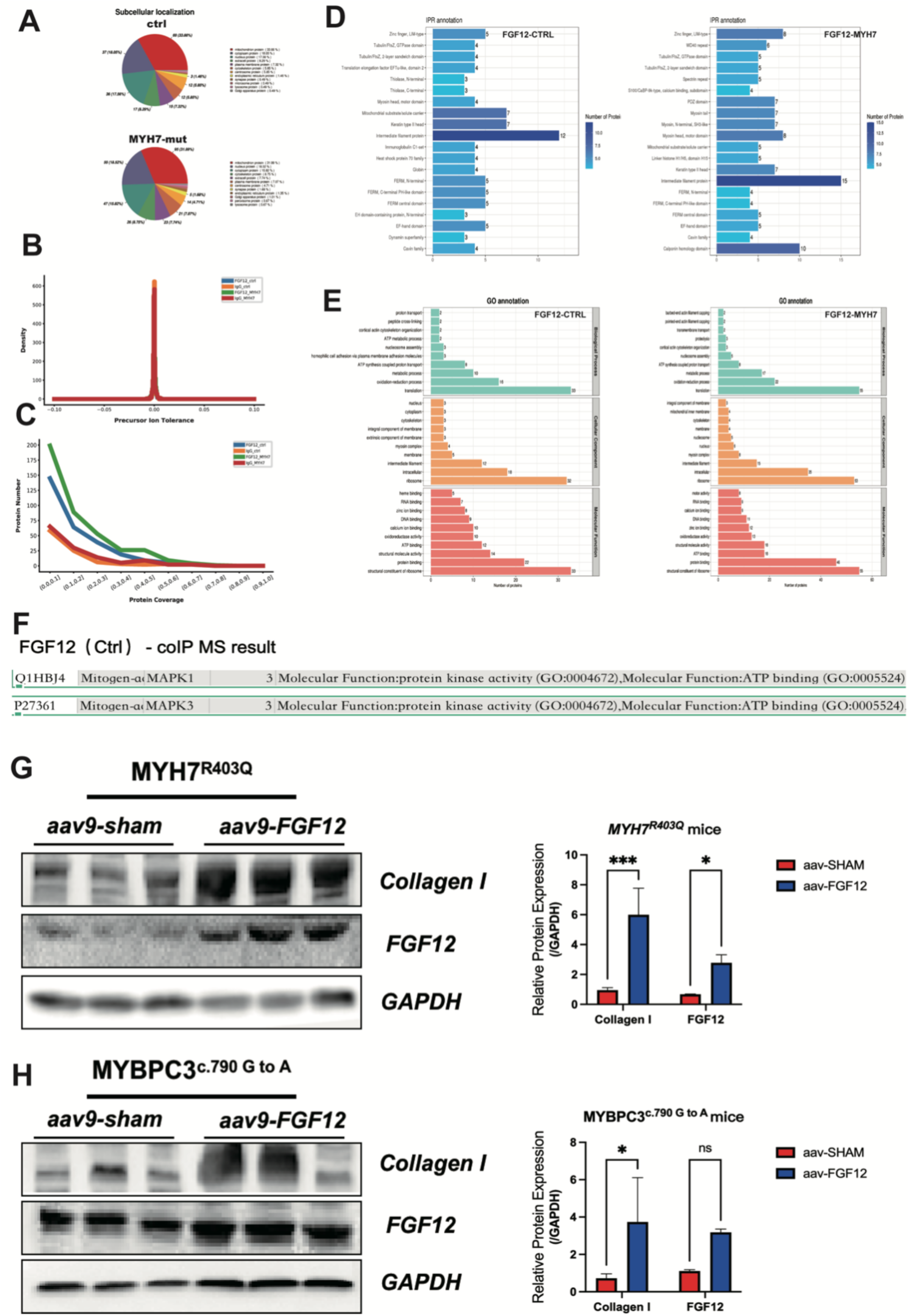
**A.** Subcellular localization of FGF12-associated binding proteins across various groups, the numbers indicate the number of protein species and percentage, respectively. Upper panel: CTRL group. Lower panel: MYH7 mutation group. **B.** Plot of the distribution of precursor ion mass tolerance across the various groups, with the horizontal axis indicating the mass deviation and the vertical axis indicating the precursor ion density distribution of the corresponding error. **C.** Plot of the coverage for the protein distribution across the various groups, with the horizontal axis representing the interval of protein coverage (length of protein covered by detected peptides/full length of given protein) and the vertical axis representing the number of proteins included across the corresponding interval. **D.** Interproscan annotation of two groups of FGF12-binding proteins, with the horizontal axis representing the number of proteins and the vertical axis representing the number of IPR entries annotated. The shade of blue represents the protein count in the FGF12-CTRL group (left) and the FGF12-MYH7 mutation group (right). **E.** The GO (Gene Ontology) annotation findings across different groups. The horizontal axis represents the number of proteins, while the vertical axis represents the GO entry annotations. The figure displays the top ten results identified in each broad category, for the FGF12-ctrl group (left) and the FGF12-MYH7 mutation group (right). **F.** MAPK1 and MAPK3 entries in the FGF12 CTRL group binding proteins. **G.** Western blot results of MYH7^R403Q^ mice using antibodies to collagen-I, FGF12, and GAPDH. The statistics of the relative expression (normalized to GAPDH) of FGF12 and collagen-I in different groups, *\*\*p <* 0.01, *\*\*\*p <* 0.005. **H.** Western blot results of MYBPC3^c.790G^ ^to^ ^A^ mice using antibodies against collagen-I, FGF12, and GAPDH. Statistics of the relative expression (normalized to GAPDH) of FGF12 and collagen-I across different groups, n.s not statistically significant \*\**p* < 0.01.

**Supplemental Figure 3.**
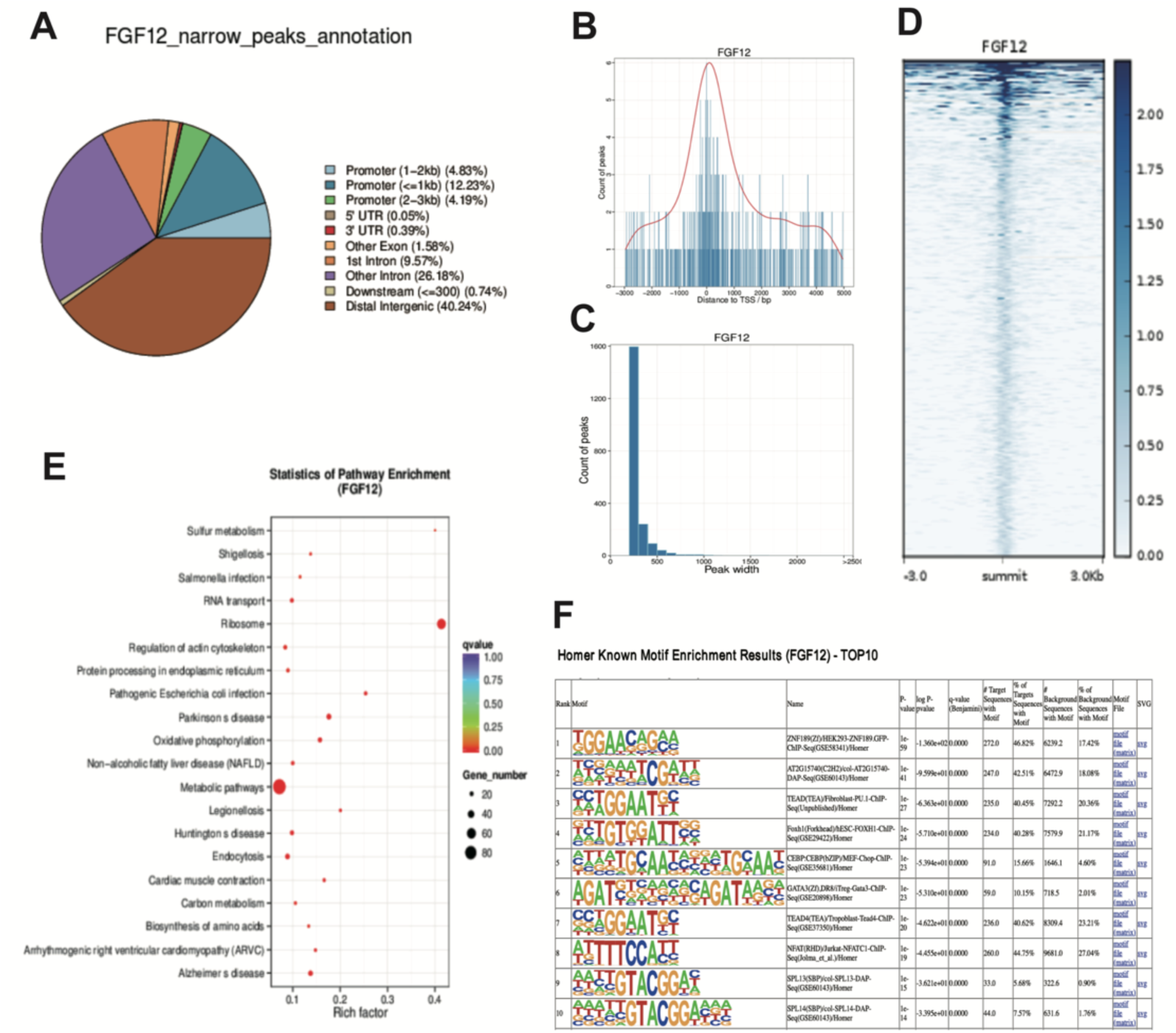
**A.** Distribution of peaks of functional regions in genes. The statistics of peaks across various functional regions of genes. **B.** Relative distribution of peaks compared to gene location. The horizontal axis displays the distance from the peak to the TSS, while the vertical axis depicts the corresponding peak count. **C.** Peak count distribution. The horizontal axis portrays the number of summits in each peak, while the vertical axis indicates the count of corresponding peaks. **D.** Reads distribution around the peak summits. A comprehensive depiction of read distribution within the 3 kb region upstream and downstream of the peak summit. The horizontal axis represents the read position relative to the peak summit, and the vertical axis showcases the clustering results of read signals in each region. **E.** KEGG enrichment scatterplot. The vertical axis denotes the pathway names, while the horizontal axis highlights the richness factor. The dot size corresponds to the quantity of peak-overlapping genes in each pathway, and the dot color signifies distinct *q*-value ranges. **F.** HOMER known motif enrichment table depicting the top ten genes.

